# Quantitative morphological variation in the developing Drosophila wing

**DOI:** 10.1101/229880

**Authors:** Alexis Matamoro-Vidal, Yunxian Huang, Isaac Salazar-Ciudad, Osamu Shimmi, David Houle

## Abstract

Quantitative variation in morphology is pervasive in all species and is the basis for the evolution of differences among species. The developmental causes of such variation are a relatively neglected research topic. Quantitative comparisons of variation arising at different developmental stages with the variation in the final structure enable us to determine when variation arises, and to generate hypotheses about the causes of that variation. We measured shape and size variation in the wing of *Drosophila melanogaster* at three developmental stages: late third instar, post-pupariation and in the adult fly. Flies of a wild-type and two mutants (*shf* and *ds*) with effects on the adult wing shape and size were studied. Despite experimental noise related to the difficulty of comparing developing structures, we found consistent differences in wing shape and size at each developmental stage between genotypes. In addition we provide linear rules allowing to link late disc morphology with early wings. Our approach provides a framework to analyze quantitative morphological variation in the developing fly wing. This framework should help to characterize the natural variation of the larval and pupal wing shape, and to measure the contribution of the processes occurring during these developmental stages to the natural variation in adult wing morphology.

## Introduction

The investigation of the developmental origins of morphological variation has become an important research area in evolutionary biology (Mallarino & Abzhanov, 2012). The complexity of developmental processes has made this investigation challenging, especially for morphological traits exhibiting multivariate and quantitative variation (Parsons & Albertson, 2013), or subtle variation at the population level (Nunes et al., 2013). While major advances have been made in finding the developmental causation of natural variation for gross morphological characteristics like the presence or absence of a structure (e.g., Arnoult et al., 2013; Chan et al., 2010), there are very few studies reporting such findings for traits exhibiting subtle and quantitative variation (Mallarino et al., 2012; Nijhout et al., 2014; Salazar-Ciudad & Jernvall, 2010).

Addressing the question of how changes in development result in quantitative variation of morphology requires quantitative comparisons of the morphology of the developing structures between individuals and between developmental stages (including the adult stage). These comparisons enable us to identify the developmental stage at which morphological variation first appears, and perhaps the developmental mechanism involved.

The wing of the fruit fly *Drosophila* is a popular model system for development and evolution. There is extensive knowledge on the variation of the adult wing shape at the intra and inter-specific levels (Houle et al., 2017). The developmental processes involved in wing shape determination are relatively well known (Diaz de la Loza & Thompson, 2016; Matamoro-Vidal et al., 2015). The fly wing goes through three main developmental stages. First, in the larval stages, the wing tissue is a mono-layered epithelium of cells, the wing imaginal disc, which undergo extensive cell division and tissue patterning. During this period, the number of cells goes from ~ 50 to ~50.000, and the major compartments of the wing (ventral, dorsal, anterior, posterior, proximal, distal) are defined. In addition, the tissue is divided into four intervein regions, separated from each other by the proveins domains which are groups of cells expressing a specific set of genes and that are the precursors of the adult wing veins L2 to L5 (Fig. 1a). Second, during metamorphosis, the wing imaginal disc is folded such that the dorsal and ventral compartments, which were on the same plane, are now apposed on each other ending up on different planes (Fig. 1b). In addition, the tissue expands in the proximo-distal axis giving the tissue a wing-like morphology (Fig. 1c). Third, during the late pupal period, a force oriented in the proximo-distal axis produced by the contraction of the hinge further elongates the tissue (Fig. 1d-e).

**Figure 1.**
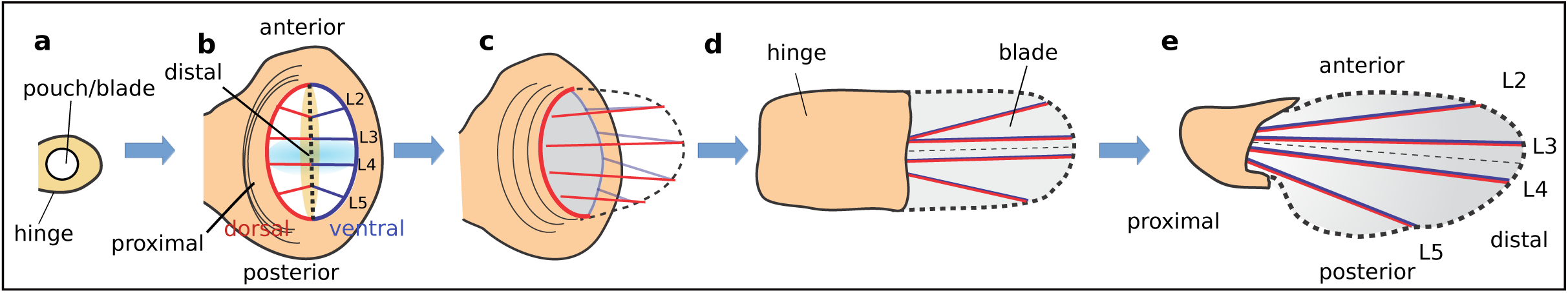
Overview of Drosophila wing development. **a.** 2^nd^ instar larval disc. **b.** 3^rd^ instar larval disc with compartments defined by the dorsal/ventral (D/V) and anterior/posterior (A/P) boundaries, provein domains (L2, L3, L4, L5) and morphogen gradients of Dpp, (produced by cells at the A/P boundary – light blue shading) and Wg, (produced by cells at the D/V boundary – orange shading). **c.** Evagination of the disc. The wing pouch folds along its D/V boundary (thick dashed line), apposing dorsal and ventral compartments, and the blade extends and become elongated along the proximal–distal axis. The part of the hinge behind the blade folds back and elongates as the blade does. **d.** Early pupal wing after evagination and expansion. **e.** Late-pupal wing. The hinge contraction creates tension that drives the elongation of the wing blade. At this stage the shape of the wing blade is similar to adult shape.

Variation in these morphogenetic events must be the source of the natural variation of the adult wing shape but the contribution of each of them is unknown. For example, the wing disc is the subject of much research in developmental biology but so far natural variation in the shape of the wing disc has not been characterized, and how changes in the shape of this structure could result in changes in the adult wing has never been investigated in a quantitative way. In this work we provide the first quantitative measurements of the developmental transformation of the late larval wing imaginal disc to the early pupal and adult wing shapes in *Drosophila melanogaster*. We compare shape variation for wing imaginal discs and early pupal wings between sexes and between three genotypes differing by mutations in two loci (*dachsous* and *shifted*) known to regulate aspects of wing development involved in the determination of adult wing shape.

## Methods

### Drosophila stocks

The number of wings examined for each condition is given on Table 1. The *yw* flies were used as wild-type. In order to compare *yw* wings with narrower wings, we studied flies homozygous for the *shf^2^* allele (Bloomington # 112), in which the spacing between the third and fourth longitudinal vein is greatly reduced (Glise et al., 2005; Gorfinkiel et al.,2005) (Figure 2). We also studied mutants of the *dachsous* (*ds*) gene, which have round wings with increased spacing between third and fourth longitudinal veins (Clark et al.,1995) (Figure 2). We used transheterozygous individuals for the alleles *ds^1^* (Bloomington # 285) and *ds^05142^* (Bloomington # 11394). A transheterozygous genotype was chosen because flies homozygous for alleles of *ds* have high lethality and severe wing overgrowth making quantitative wing shape measurements challenging. *ds^1^* and *ds^05142^* lines were balanced over the Cyo, Dfd-YFP balancer chromosome and crossed with each other. *ds^1^/ds^05142^* flies were thus identified by lack of YFP.

**Figure 2.**
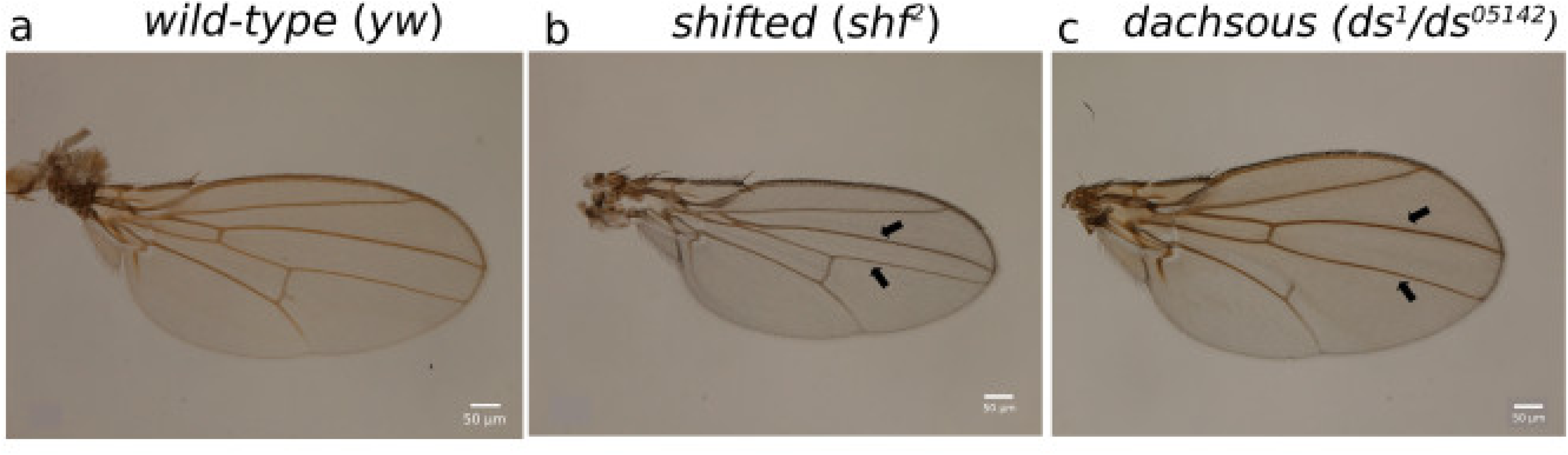
Adult wings for the three genotypes studied. **a**. *yw*. **b.** *shf^2^*. **c.** *ds* (*ds^1^/ds^05142^*). Black arrows highlight the longitudinal veins 3 and 4.

**Table 1.** Sample size, area means and standard deviations by stage and genotype.

|  | genotype | N | Area (mm<sup>2</sup>) | Areas as proportions of total |  |  |
| --- | --- | --- | --- | --- | --- | --- |
|  |  |  |  | Anterior | Middle | Posterior |
| Larval | *yw* | 16 | 0.0100±0.0015 | 0.547±0.042 | 0.165±0.018 | 0.288±0.030 |
|  | *ds* | 8 | 0.0102±0.0028 | 0.560±0.029 | 0.137±0.025 | 0.304±0.014 |
|  | *shf2* | 9 | 0.0077±0.0008 | 0.558±0.030 | 0.120±0.009 | 0.322±0.032 |
| Pupal | *yw* | 15 | 0.0243±0.0041 | 0.530±0.043 | 0.143±0.023 | 0.327±0.042 |
|  | *ds* | 12 | 0.0210±0.0042 | 0.536±0.034 | 0.156±0.020 | 0.308±0.026 |
|  | *shf2* | 15 | 0.0182±0.0049 | 0.613±0.043 | 0.080±0.013 | 0.308±0.035 |
| Adult | *yw* | 6 | 0.0976±0.0099 | 0.452±0.010 | 0.166±0.014 | 0.382±0.009 |
|  | *ds* | 12 | 0.1286±0.0140 | 0.446±0.006 | 0.176±0.004 | 0.378±0.006 |
|  | *shf2* | 16 | 0.0907±0.0097 | 0.495±0.008 | 0.116±0.009 | 0.389±0.012 |

**Table 2.** Results from MANOVA of shape data within each stage.

| Stage | Effect | Num df | den df | Wilks' $\lambda$ | P |
| --- | --- | --- | --- | --- | --- |
| Larva | Genotype | 36 | 22 | 0.004 | <0.0001 |
|  | sex | 18 | 11 | 0.208 | 0.08 |
|  | Genotype by sex |  |  |  | >0.2 |
| Pupa | Genotype | 36 | 38 | 0.030 | <0.0001 |
|  | sex | 18 | 19 | 0.417 | 0.20 |
|  | Genotype by sex | 36 | 38 | 0.177 | 0.13 |
| Adult | Genotype | 36 | 22 | 0.0002 | <0.0001 |
|  | sex | 18 | 11 | 0.207 | 0.08 |
|  | Genotype by sex | 36 | 22 | 0.082 | 0.15 |

### Dissections

Larval wing discs were dissected from wandering third instar larvae. The wing discs were fixed with 4% Formaldehyde fixative at room temperature for 20mins, then dissected from the larva.

Pupal wings were dissected from pupae aged from the white prepupal stage. White prepuape were defined as individuals that had ceased movement, everted anterior spiracles, but had not yet begun tanning of cuticle. Individual white prepuape were picked and reared at 25°C until dissection. The pupal wings were fixed with 4% Formaldehyde fixative at 5 h after pupariation, left at at 4 °C overnight, and then dissected from the pupae.

Adult wings were dissected from adult flies and mounted with 80% glycerol.

### Immunostaining

We used immunological stains to identify the positions of proveins in larval wing discs and pupal wings. Immunostaining was performed as previously described (Matsuda et al 2013). The primary antibodies used were mouse anti-Delta at 1:50 (Developmental Studies Hybridoma Bank (DSHB), rat anti-cubitus at 1:50 (DSHB). The secondary antibodies were as follows: goat anti-mouse IgG-Alexa 568 and goat anti-rat IgG-Alexa 488 were used at 1:200, respectively (Invitrogen).

### Imaging

The fluorescent images were obtained with Zeiss LSM700 confocal microscope. Adult wing images were obtained with Nikon eclipse 90i.

### Landmarks and semi-landmarks

Size and shape of 3^rd^ instar wing discs, 5 h pupal wings, and adult wings were measured by gathering a set of 8 landmarks and 9 semi-landmarks on each specimen (Figure 3), using *tpsUtil* and *tpsDig2* software (http://life.bio.sunysb.edu/morph) for the discs and pupal wings; and using *Wings4* (Houle et al., 2003; http://bio.fsu.edu/˜dhoule/wings.html) for the adult wings.

**Figure 3.**
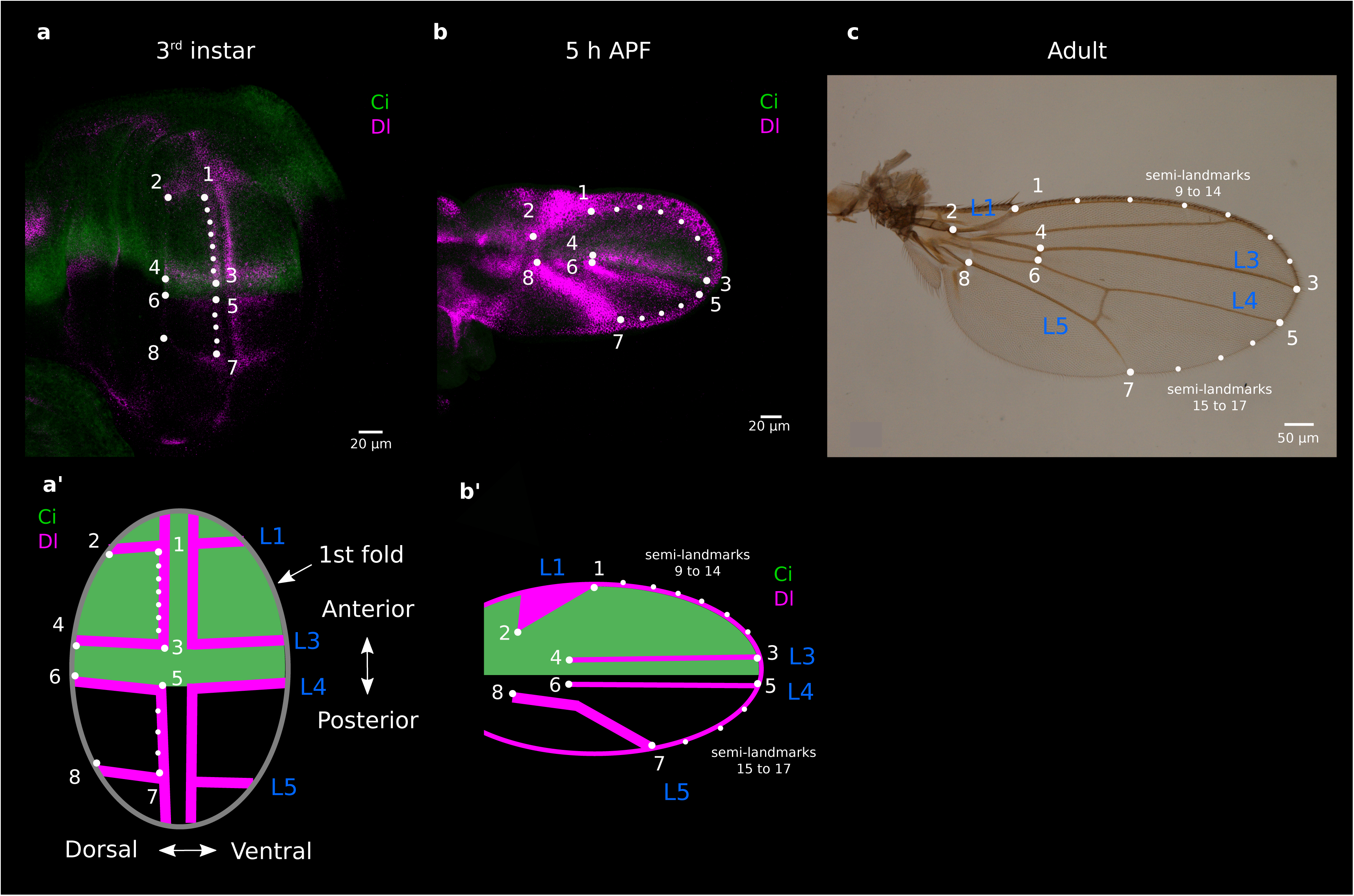
Landmarks and semi-landmarks used for the morphometric analyses. **a**. 3^rd^ instar larval wing stained with antibodies against Cubitus interruptus (Ci, green) and against Delta (Dl, magenta). Wing shape was measured by gathering 8 landmarks (big white dots numbered 1-8) and 9 semi-landmarks (smaller white dots). **a'.** Diagram of a 3^rd^ instar larval wing showing how the Delta staining (proveins and D/V boundary) and the 1^st^ fold were used for landmarks and semi-landmarks positioning. **b.** Pupal wing at 5 h after puparium formation (APF) with same staining than in 'a' and landmarks/semi-landmarks positions hypothesized to be homologous to those in 'a'. **b'.** Diagram of 5 h APF pupal wing showing how Delta staining (proveins), and the wing margin were used for landmarks and semi-landmarks positioning. **c.** Dorsal adult wing with landmarks and semi-landmarks positions hypothesized to be the same than in 'a' and 'b'.

The positions of the landmarks were defined using molecular and morphological markers (Figure 3). For the former, we used immunostaining showing the Cubitus interruptus (Ci) and Delta (Dl) territories in wing discs and 5h pupal wings. The gene *ci* is expressed in all the anterior wing whereas *dl* is expressed in two stripes of cells following the dorso-ventral boundary, as well as in the proveins territories precursors of the veins 1, 3, 4 and 5 (Biehs et al., 1998; Cook et al., 2004). The morphological markers were the 1^st^ fold of the wing pouch, the margins of the pupal and adult wings, and the veins of the adult wings.

For the wing discs, four landmarks (1, 3, 5 and 7) were placed in the distal part of the tissue, at the intersections of the DV boundary with the proveins L1, L3, L4 and L5, respectively. Four other landmarks (2, 4, 6 and 8) were placed at the distal tips of the proveins 1, 3, 4 and 5, respectively, which coincide with the intersections of these proveins and the 1^st^ fold of the pouch. Note that the position of vein L4 coincides with the end of the anterior compartment (shown by Ci territory). In addition, two sets of semi-landmarks were placed on the DV boundary. The first one (9-14) was placed in the portion of the DV boundary contained within proveins L1 and L3, and the second one (15-17) was placed in the portion within L4 and L5. Data were initially collected for ventral and dorsal compartments of the wing disc. However, the ventral compartment was found to be quite variable because this part of the disc starts to evert very early. Thus only the data for the dorsal disc were considered.

For the pupal wings, four landmarks (1, 3, 5 and 7) were placed at the intersections of proveins L1, L3, L4 and L5 with the wing margin, and four others (2, 4, 6 and 8) at the proximal tips of proveins L1, L3, L4 and L5. As in the wing discs, two sets of semi-landmarks were placed along the wing margin. One (9-14) was placed in the portion of the wing margin contained within proveins L1 and L3, and another (15-17) in the portion within L4 and L5.

For the adult wings, four landmarks (1, 3, 5 and 7) were placed at the intersections of veins L1, L3, L4 and L5 with the wing margin. Landmarks 4 and 6 were placed at the intersections between the anterior cross-vein and veins L3-L4; landmark 8 was placed at the intersection between veins L5 and L6 (anal crossvein) and landmark 2 at the proximal end of vein L1. Again, two sets of semi-landmarks were placed along the wing margin between L1-L3 (9-14) and L4-L5 (15-17).

### Shape analysis

The combined data on landmark and semi-landmark positions from the larval discs and the pupal and adult wings was subjected to generalized Procrustes superimposition (Rohlf & Slice, 1990), using the program tpsRelw (http://life.bio.sunysb.edu/morph/index.html). Procrustes superimposition scales forms to the same size, translates their centroids to the same location, and rotates them to minimize the squared deviations around each point. This separates the useful size and shape information from the nuisance parameters introduced by the arbitrary location and rotation of the specimens within the images. The positions of the semi-landmarks were slid along each dorsal-ventral boundary segment defined by the boundary landmarks to minimize deviation along the segment using the standard model in tpsRelw (Rohlf). Although we measured the x and y coordinates of 17 landmarks and semi-landmarks, there were only 18 degrees of freedom in the shape data after registration and sliding.

Analysis of shapes using tpsSmall (http://life.bio.sunysb.edu/morph/index.html) shows that Euclidean distances where extremely highly correlated with Procrustes distances (r=0.999964), despite the wide differences in shapes of larval, pupal and adult forms. We performed a principal component analysis on the shape data, retaining 18 PC axes for further analyses.

Outliers were diagnosed using a robust approach for the first 5 shape principal component axes within each genotype and stage using the Diagnostics option in the Robsutreg procedure in SAS, employing a dummy dependent variable. Specimens more than 3 S.D.s away from the robust means were identified as outliers. Images of putative outliers were re-examined to determine the source of the unusual measurements. For adult wings, wings with relatively extreme ds and shf2 phenotypes were identified by Robustreg as outliers. We retained these in the data, as the deviations were relatively modest. For larval wings, one shf2 outlier appeared to have a damaged disc, and was omitted. Four pupal outliers (two ds, and two yw) greater than 6 S.D. from the robust mean proved to have unusual staining patterns, or distortions of the epithelia, and these were omitted. The final shape data set consists of 108 specimens. No univariate outliers for size (area or centroid size) were detected using Grubb’s test.

To test whether genotypes differed in the developmental transformations they undergo from larval to pupal to adult form, we used a multivariate analysis of variance (MANOVA). Type III sums of squares and cross-products were used to calculate test statistics. The variance of shape was very different among developmental stages, which violates the assumption of homogeneous variances used for conventional statistical tests. To provide an alternative test, we performed MANOVAs of data randomized to make the null hypothesis of no effect true. We first decomposed each observation into the grand mean, plus residuals corresponding to stage, genotype, and genotype by stage data, and residual as follows

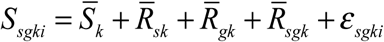

where *s* indexes developmental stage, *g* indexes genotype, *k* indexes the shape variable, i the individual, the overbar indicates a mean shape, and ^*ε*^^*_sgki_*^ is the deviation of the individual from the stage-genotype mean. We then randomized just the deviations used to test a particular hypothesis, holding all other aspects of the observation constant. For example, to test for stage by genotype interactions, we randomized 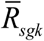 values among individuals within stages. The values of Wilks’ lambda were retained from 1,000 randomized analyses, and compared with the Wilks’ lambda obtained from analyzing the observed data.

### Scalar measures

The standardized distances between the 28 possible pairwise combinations of the 8 landmarks were obtained from the Cartesian coordinates of the landmarks corrected by the centroid size. Centroid size is proportional to the square root of wing area. In addition, we measured the standardized lengths for a portion of the anterior margin using landmarks 1-3 and semi-landmarks 9-14, and for a portion of the posterior margin using landmarks 5-7 and semi-landmarks 15-17 (Figure 4).

**Figure 4.**
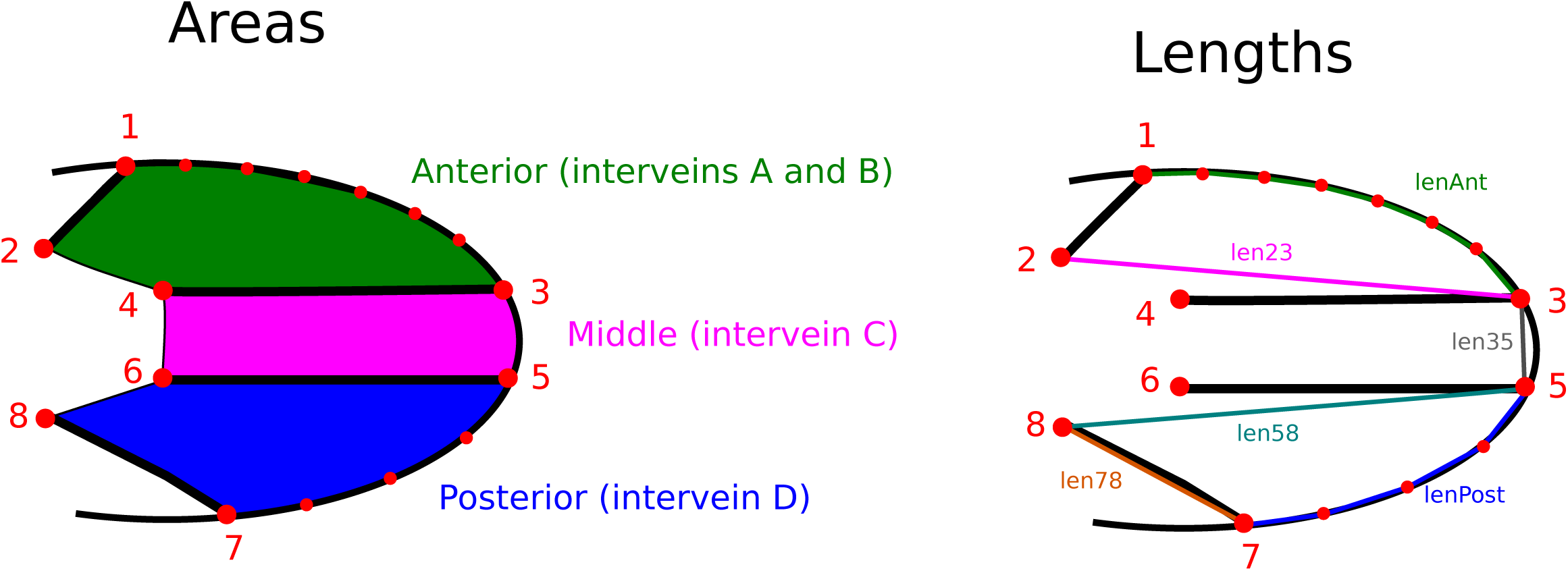
Wing diagrams illustrating areas and lengths compared over stages and genotypes. Three areas were measured: interveins A and B (anterior, green), intervein c (middle, magenta) and intervein d (posterior, blue). We measured the distances between the twenty-eight possible pairwise combinations of the eight landmarks. Only four distances are shown in the diagram for clarity, between landmarks 2-3 (len23); 3-5 (len35); 5-8 (len58) and 7-8 (len78). In addition, we measured the length of a portion of the anterior margin (lenAnt) and of a portion of the posterior margin (lenPost) using landmarks landmarks 1, 3, 5, 7 and the semi-landmarks.,

Three areas were calculated using the surveyor’s formula for calculating areas of polygons. The first area was obtained by calculating the area of the regular polygon within the landmarks 1-4 and semi-landmarks 9-14, thus obtaining a proxy of the anterior wing area (‘Anterior’). The second areas is for the polygon defined by the landmarks 3-6 which contains the region within the longitudinal veins L3 and L4 (‘Middle’). The third area is the one of the regular polygon defined by landmarks 5-8 and semi-landmarks 15-17, and gives a proxy of the posterior wing area (‘Posterior’) (Figure 4).

Standardized lengths and areas were compared between developmental stages and between genotypes by calculating means ratios. Values of variance for these ratios were obtained by bootstrapping the data. For example, change in the standardized length between landmarks 2 and 8 (stlen28) during the larval to pupal transition in the *yw* genotype was calculated with the following procedure: individual values for stlen28 in the *yw* pupal wings population were re-sampled with replacement a number of times equal to the number of individuals in the population. The mean on the re-sampled data was calculated and divided by the mean obtained by the same approach on the *yw* larval wing population. This procedure was repeated 1000 times providing thus a distribution of values for the ratio of stlen28 (pupa) / stlen28(larva) of the *yw* genotype.

### Analyses

Statistical tests were carried out in the GLM procedure in SAS (), assuming that stage, genotype and sex are fixed factors. Type III sums of squares and cross-products were used for statistical testing. When interaction terms had P>0.2, they were dropped from the final model. Post-hoc comparisons among genotypes were adjusted within traits for multiple comparisons using the Tukey-Kramer method. The standard errors of ratios of wing areas were approximated using standard formulas for the variance of a ratio, and tests for differences among ratios assumed that the differences are normally distributed. To do this formula, we had to assume that the covariance of areas between stages is 0, leading to an overestimate of the variance, and conservative tests for differences among the ratios.

## Results

### Size over stages

Means and standard errors for areas are shown in Table 1. Analysis of log10 area in a model with stage, sex and genotype as factors shows that the between stage differences are highly significantly different from 0 (P<0.0001 for all comparisons).

Ratios for changes in area between stages are shown in suppl. figure 1. The dorsal area expands markedly during development. The ratios of pupal to larval wing areas are similar for the three genotypes, increasing by factors of 2.1 ± 0.2 (*ds*), 2.3 ± 0.2 (*shf2*) and 2.4 ± 0.1 for the wild-type (*yw*). At the pupal to adult transition the increases in wing area are all significantly different from each other, increasing by factors of 6.1 ± 0.4 for *ds*, 5.0 ± 0.4 for *shf2*, and 4.0 ± 0.2 for *yw*. Wing area increases between the larval and adult stages by factors of 12.6 ± 1.3 (*ds*), 11.8 ± 0.6 (*shf2*) and 9.7 ± 0.5 (*yw*). The growth ratio of *yw* is significantly lower than those of the other two genotypes.

### Shape over stages

To examine the relative shapes of individuals at each stage, we performed canonical discriminant analyses on the principal components of the shape data. Figure 5 plots the scores on the first and second canonical axes when the discriminant analysis used developmental stage as the classification variable. Larval, pupal and adult shapes are extremely distinct. Note that the variation among individuals within stages is quite different. As a result, standard statistical tests across stages are likely to be biased. A MANOVA on the shape data showed that the effect of stage was highly significant (Wilks’ λ=0.00159, num df=38, den df=158, P<0.0001).

**Figure 5.**
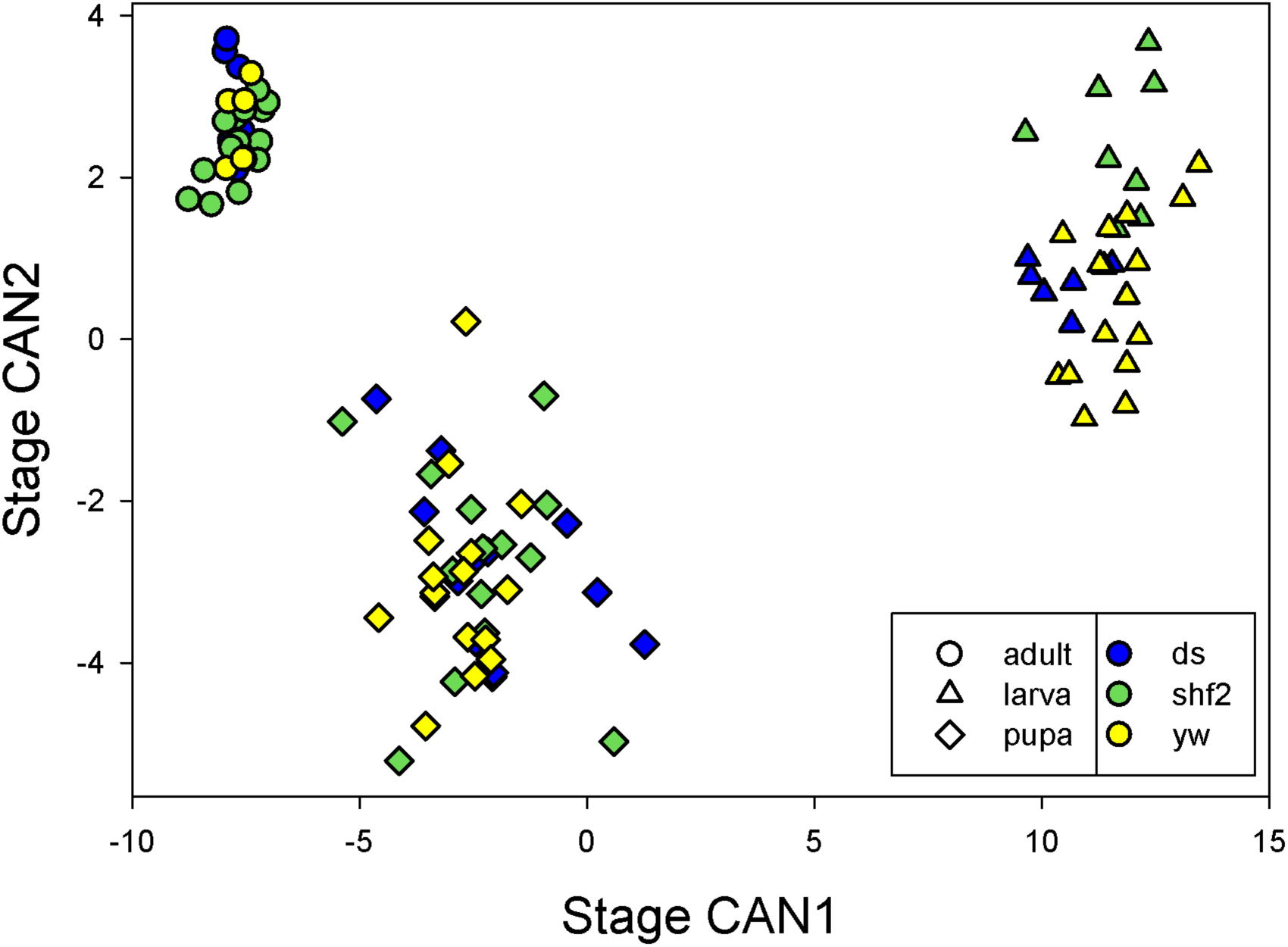
Scores for shape on canonical axes chosen to discriminate stages.

To enable visualization of shape differences, we used the program Lory (Márquez et al., 2012) to show one pattern of relative expansion or contraction that can transform one mean shape into another. Figure 6 plots stage transformations. The magenta arrows represent changes in relative locations of landmarks, while the colors between landmarks represent the inferred expansion and contractions that can bring about the changes in landmark positions. It is important to realize that these represent only shape change, and not size change. The transformation shown is a hypothesis, as other patterns of expansion and contraction can lead to the same shape change at the measured locations.

**Figure 6.**
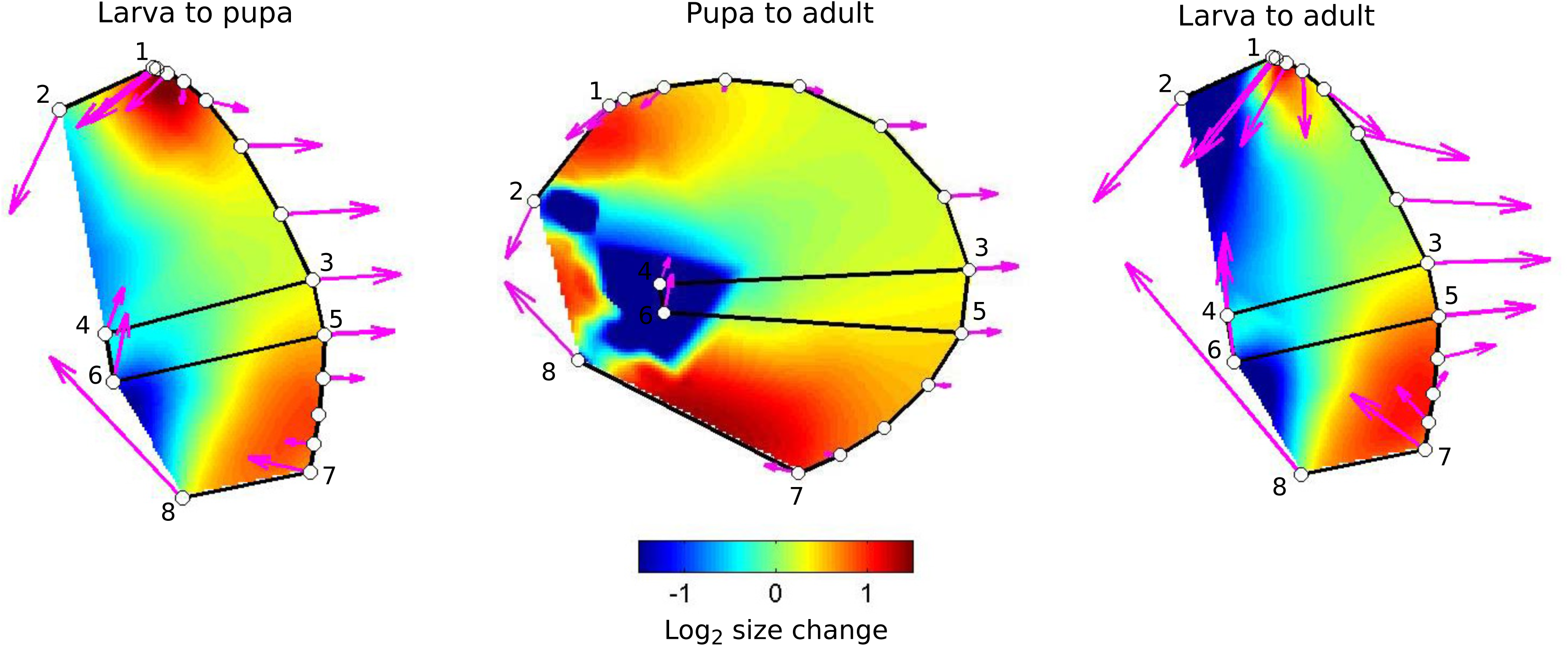
Differences among stages. Colors represent inferred changes in the relative areas of parts of wing necessary to transform the form from the earlier stage (e.g. larva) to the later (e.g. adult) stage. Expansions and contractions are shown on a log2 scale, the orange at +1 represents a doubling to relative area, while blue at −1 represents a local halving of area. Magenta arrows represent the pattern of change in location of landmarks (numbered 1 to 8) and of semi-landmarks.

The overall pattern of shape change is that the distal part of the wing, closest to veins L3 and L4, move to the right in the figure, shown by the magenta arrows, while the proximal anterior and posterior parts of the boundary are drawn together and to the left, relative to the rest of the wing. Movie 1 shows the same data as a transformation of the outline of the wing between stages.

Our linear measurements show that during the larval to pupal phase, shape change is characterized by a narrowing of the tissue along the anterior-posterior axis (e.g., reduction in the relative distances between pairs of landmarks 2-8; 1-8; 4-8 – suppl. figure 2a) and by an expansion in the direction of the proximal – distal axis, as illustrated by the increase in the relative distances between the pairs of landmarks 7-8; 5-8; 3-8, and by the lengthening of the anterior and posterior margins (suppl. figure 2a). This pattern of shape change is continued into late pupal development, with a pronounced constriction along the anterior-posterior axis in the proximal parts of the wing (~ 50 % decrease in the distance between the pairs of landmarks 2-8;1-4;1-8 and 1-6) and elongation along the proximal-distal axis (suppl. figure 2B).

### Differences in shape among genotypes

We tested for differences in shape between genotypes within stages using a multivariate analysis of variance, with the results shown in Table 3. In all three stages, there were highly significant differences among genotypes.

**Table 3.** Mean shape distance between individuals in each stage/genotype combination. Values are the mean Euclidean distances between the 34 element vector of shape coordinates. Diagonals are the average distances between different individuals of the same genotype and stage.

|  |  | Larva |  |  | Pupa |  |  | Adult |  |  |
| --- | --- | --- | --- | --- | --- | --- | --- | --- | --- | --- |
|  |  | *yw* | *ds* | *shf2* | *yw* | *ds* | *shf2* | *yw* | *ds* | *shf2* |
| Larva | *yw* | 0.12 | 0.14 | 0.16 | 0.56 | 0.50 | 0.50 | 0.77 | 0.75 | 0.77 |
|  | *ds* |  | 0.10 | 0.20 | 0.54 | 0.47 | 0.48 | 0.75 | 0.73 | 0.76 |
|  | *shf2* |  |  | 0.10 | 0.54 | 0.50 | 0.48 | 0.73 | 0.72 | 0.73 |
| Pupa | *yw* |  |  |  | 0.15 | 0.21 | 0.21 | 0.28 | 0.24 | 0.30 |
|  | *ds* |  |  |  |  | 0.17 | 0.22 | 0.36 | 0.31 | 0.38 |
|  | *shf2* |  |  |  |  |  | 0.18 | 0.34 | 0.32 | 0.35 |
| Adult | *yw* |  |  |  |  |  |  | 0.02 | 0.11 | 0.07 |
|  | *ds* |  |  |  |  |  |  |  | 0.04 | 0.16 |
|  | *shf2* |  |  |  |  |  |  |  |  | 0.02 |

Figure 7 plots the differences between genotypes relative to the *yw* genotype. We used *yw* as the reference as the mutations it carries are not known affect wing development. Comparison of *yw* and *ds* suggest that differences in the anterior- and posterior-most regions that will become proximal in the adult exist from the larval stage, but that the majority of the difference between these genotypes arise during pupal development, and the peripheral areas of the blade expand more in ds mutants than *yw.* Comparison of *yw* and *shf2* suggests that the region between L3 and L4 is markedly smaller in *shf2* from the larval stage. This contraction persists, but is balanced principally by an expansion of the proximal part of the wing anterior to L3 in later stages.

**Figure 7.**
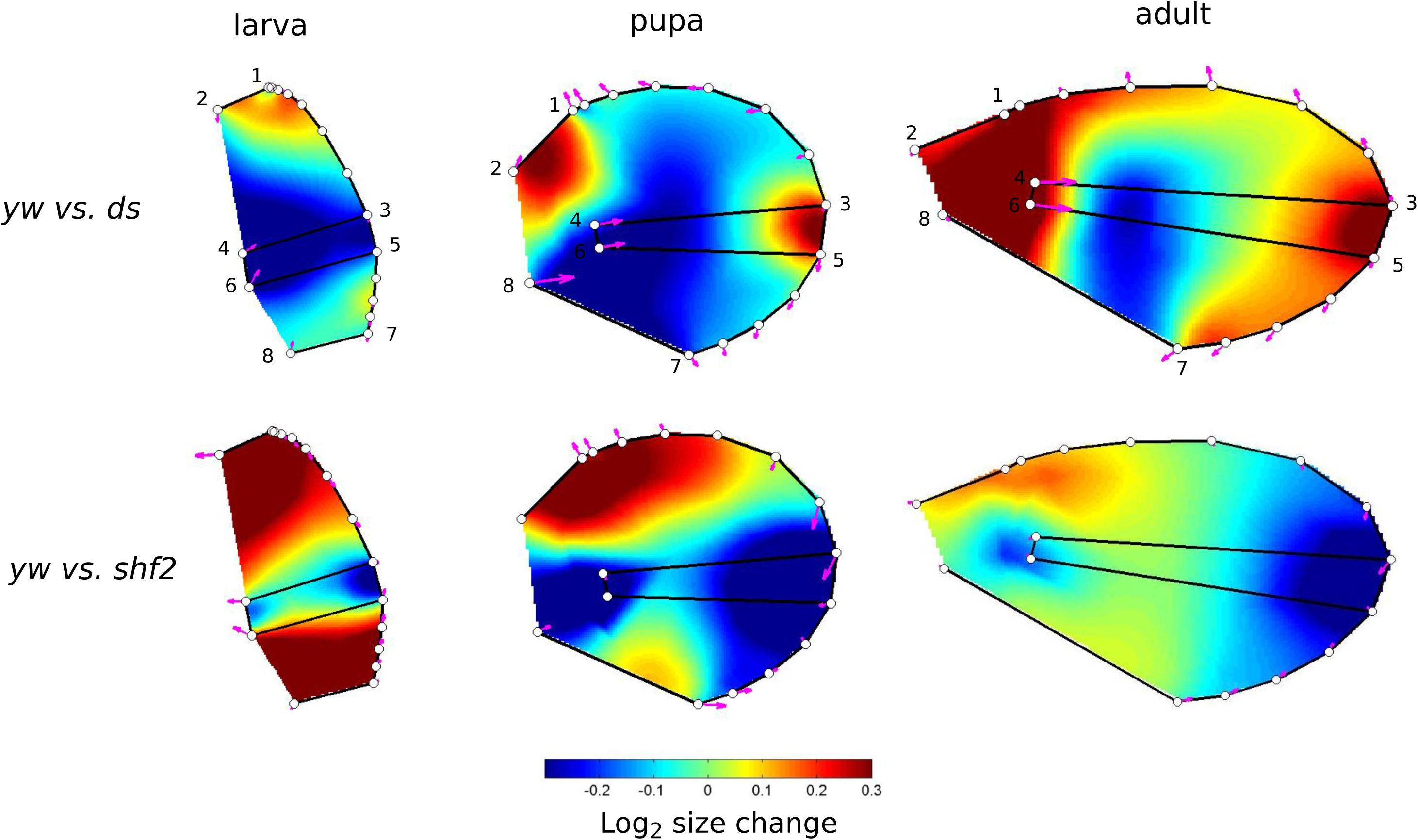
Differences among genotypes within stages. The *yw* genotype is taken as the reference, and colors represent changes in relative area necessary to transform the wing at a given stage (larva, pupa or adult) into the other two genotypes. Note that the scale differs from that in Fig. 6. The top of the scale represents an increase by a factor of 1.23.

To diagnose where these differences arise we examined the ratios *ds/yw* and *shf2/yw* of the standardized lengths and areas. These ratios were first conducted on the adult data to see what is different in adult wings between *yw* and the mutants, and then on the larval and pupal wing data to check when the variation observed in the adults appears during development.

The ratios *ds/yw* of the standardized lengths for the adult wings are shown in suppl. fig. 3A. The *ds* adult wings are narrower relative to *yw* along the P/D axis in the distal part, as well as broader along this same axis in the proximal part. This is shown by the shift of landmarks 4 and 6 towards the distal parts. These two landmarks indeed have higher relative distances with respect to landmarks 1, 2 and 8, as well as lower relative distances with respect to landmarks 3, 5 and 7. In addition, *ds* wings are broader along the anterior-posterior axis, as shown by increase in relative distances between the pairs of landmarks 4-6 and 3-5. Regarding the areas (suppl fig 3B), our data show that *ds* wings are 1.3 times bigger than *yw*, and this is due to an increase in all the three areas measured with a slightly more important contribution of the “Middle” area.

Examining these ratios in the larval and pupal wings shows that the differences observed between *ds* and *yw* adult wings appear at different times during development. The proximo-distal narrowing of the distal part of the wing is observed in the larval stage (Figure 8a, suppl. fig. 4a), whereas the proximo-distal lengthening of the proximal wing, as well as the broadening in the A/P axis appears at the pupal stage (Figure 8b, suppl. fig. 4b). The variation in wing area occurs mostly during the pupal to adult transition, as well as the shift of landmarks 4-6 towards the distal parts of the wing (Figure 8c, suppl. fig. 4c).

**Figure 8.**
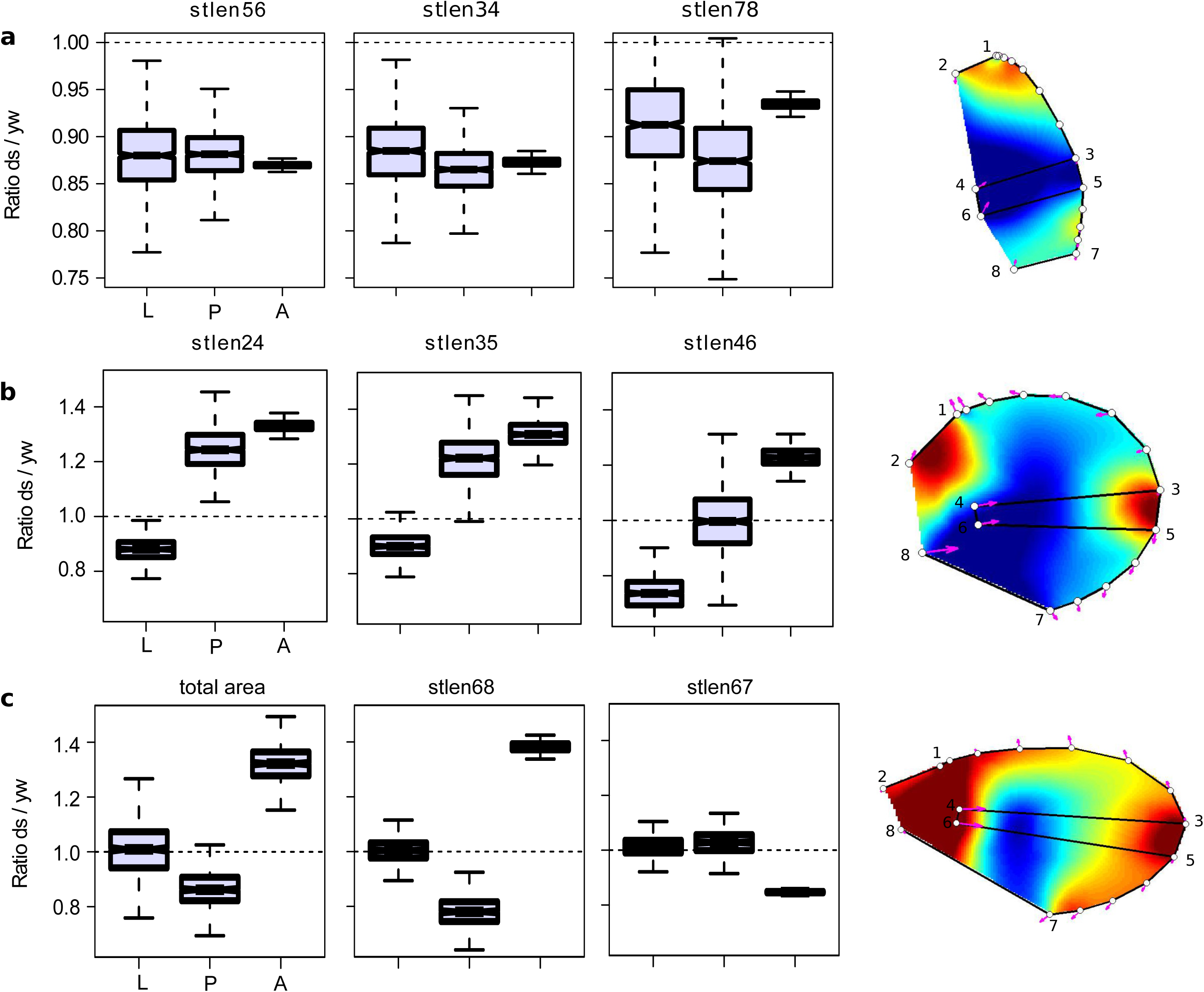
Developmental stage at which the adult wing shape differences between *ds* and *yw* appear. The boxplots show ratios of means between *yw* and *ds* genotypes at each stage for standardized distances between the pairs of landmarks (stlen) and total wing area (total area). Variance for the ratios were obtained by bootstrap (n = 1000, see methods). Notches on the boxplots display the 95 % confidence interval around the median. For clarity, only few representative variables are shown (see suppl. figure 4 for the other variables). **a.** Variables for which the differences between *ds* and *yw* adult wings appear before the 3^rd^ instar larval stage. **b.** Variables for which the differences between *ds* and *yw* adult wings appear during the larva to pupa transition. Note that for stlen46, there is a continuous increase of the ratio during larval and pupal development to reach the adult ratio. **c.** Variables for which the differences between *ds* and *yw* adult wings appear during the pupa tu adult transition. L, larva; P, pupa; A, adult. For each stage, a diagram showing the overall shape difference between genotypes (from Fig. 7) in shown.

Supplementary Figure 5a shows the ratios *shf2/yw* for the standardized lengths in adult wings. The principal differences are the reduction of the distances between the pairs of landmarks 3-5 and 4-6, in *shf2*, and the corresponding reduction in that area of the wing (intervein L3L4) (suppl. fig 5b). In the case of *shf2,* the differences observed in the adult wings are established in the larval wing (Figure 9), with relatively small changes after that stage.

**Figure 9.**
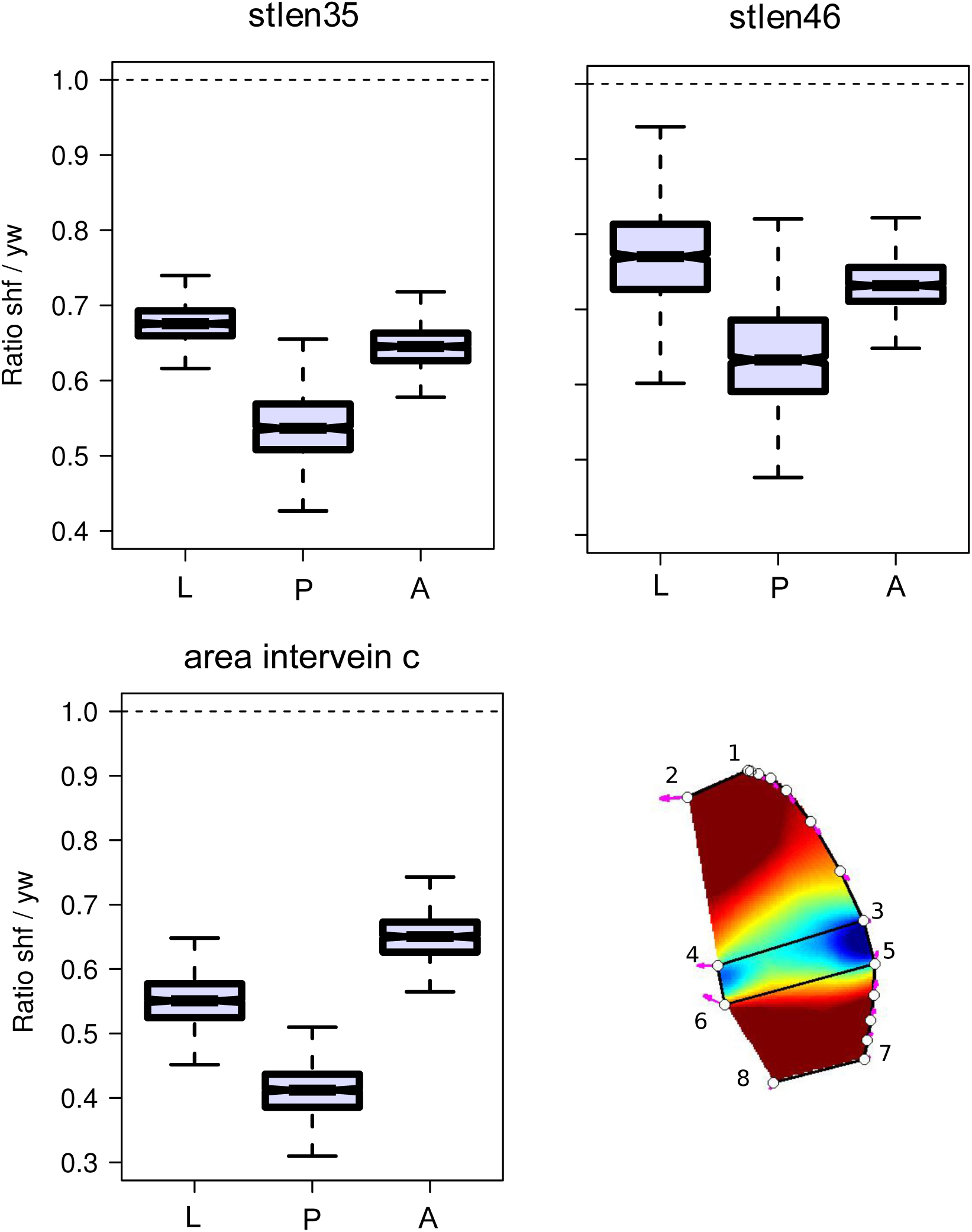
Developmental stage at which the adult wing shape differences between *shf2* and *yw* appear. The boxplots were obtained as in Figure 8. All major differences between *shf2* and *yw* adult wings are observed since the 3^rd^ instar larval stage. L, larva; P, pupa; A, adult. A diagram showing the overall larval wing shape difference between genotypes *yw* and *shf2* (from Fig. 7) is shown.

### Shape transformations between stages

Randomized MANOVA analysis showed a highly significant effect of genotype over stages (Wilks’ λ=0.078, minimum of 1,000 randomized Wilks’ λ=0.244). This result demonstrates that some of the differences among genotypes are consistent across all three stages. There was also a highly significant stage by genotype interaction (Wilks’ λ=0.078, minimum of 1,000 randomized Wilks’ λ=0.568), which demonstrates that there are changes in the relationships among genotypes over stages.

Figure 10 shows the scores on the first and second canonical axes when the discriminant analysis used genotype as the classification variable. Genotypes are well separated on these axes, with a few exceptions. The similar locations of genotypes across stages suggests that shape differences in the larva are retained through the pupa and adult shapes.

**Figure 10.**
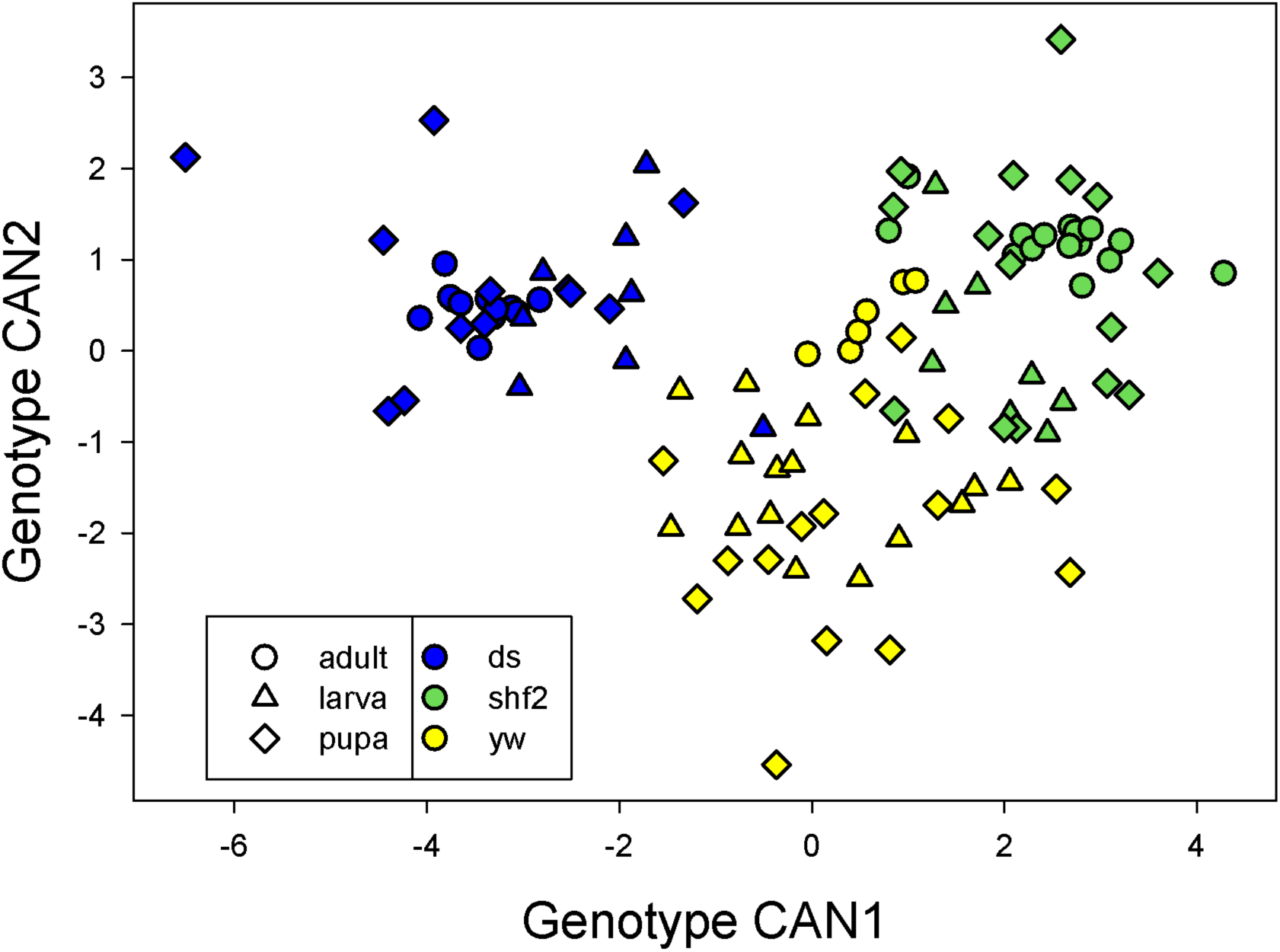
Scores for shape on canonical axes chosen to discriminate genotypes.

To get a sense for the size of stage and genotype effects, we calculated the matrix of Euclidean distances in shape space (centroid size units) among individuals in each stage/genotype combination, with the results shown in Table 3. The mean distance between individuals within stages is 0.13 (0.14 within larvae, 0.19 with pupae, 0.07 within adults,), while it is 0.51 between larval and pupal shapes, 0.32 between pupal and adult shapes, and 0.74 between larval and adult shapes. Thus, pupal shape is more similar to adult shape than to larval shape, suggesting that eversion and folding has a larger effect on shape than pupal development. The differences in shape among genotypes within stages are less dramatic. For larvae the average distance between different individuals with the same genotype is 0.11, while the differences among individuals of different genotypes is 0.17. In pupae the within genotype distances average 0.17, while the among genotype distances average 0.22. Adults of the same genotype average just 0.03 in distance, while the among genotype distances average 0.11. This is likely to be due to higher accuracy of measurements in adults.

The genotype-stage interactions demonstrate that the developmental transformations between stages differ among genotypes. To get a sense for the magnitude of the genotype-stage interactions, we calculated the angles between shape change vectors. To do this, we calculated the average direction of shape change between stages for each genotype as the difference in mean phenotype across each transition. We then calculated the angles between these shape change vectors, with the results shown in Table 4. Completely independent shape changes would have an angle of 90, while identical transformations have an angle of 0. The angles are quite close to the minimum of 0, and suggest that genotypes are undergoing similar transformations. In particular, the transformations from larval to adult shapes differ on average by just 8 degrees. Angles involving pupal shapes are generally larger., which probably reflects the larger variation in pupal shape than the other two stages, with correspondingly larger uncertainty as to the true pupal mean.

**Table 4.** Angle in degrees between the vectors of shape changes for each genotype.

| Comparison | *yw* vs. *ds* | *yw* vs. *shf2* | *ds* vs. *shf2* |
| --- | --- | --- | --- |
| larval to pupal | 9.9 | 16.0 | 16.8 |
| pupal to adult | 25.4 | 14.3 | 19.4 |
| larval to adult | 7.8 | 6.3 | 9.8 |

## Discussion

Attempts to understand the developmental causes of quantitative natural variation have been hindered by the complexity of the developmental processes (Parsons & Albertson, 2013). Even in a relatively simple structure such as the Drosophila wing, many developmental processes contribute to morphogenesis (Matamoro-Vidal et al., 2015). The tremendous progress of developmental biology in quantifying many aspects of morphogenesis makes it likely that these difficulties can be overcome (Oates et al., 2009). We have used a framework based on geometric morphometrics that allows us to quantify wing shape and size variation during development. We have applied this to determine when genotypic differences in wing shape and size appear during development. By narrowing the time frame when differences arise, we can narrow down the number of candidate developmental processes that potentially cause genotypic differences.

In the language of geometric morphometrics, features sharing identity or homology among specimens are referred to as landmarks. We used the relative positions of landmarks to compare changes in size and shape between developmental stages and genotypes. Landmark homology is assured when comparing wing specimens at the same developmental stage, but is not always clear when comparing specimens at different developmental stages. These uncertainties urge some caution in interpreting our results. For example, the dorsal-ventral (D-V) boundary in the larval disc is undoubtedly homologous with the wing margin in pupae and adults. On the other hand, the position of landmarks along the D-V boundary defined by Delta expression (landmarks 1, 3, 5 and 7) may be shifted along that boundary relative to those visible in the adult wing. The homology of the other four landmarks (2, 4, 6 and 8) is less assured across developmental stages, particularly when compared to the adult wing. However, it seems likely that discrepancies in the placement of these landmarks will be consistent among genotypes. If this assumption is met, differences among genotypes (and sexes) in how these landmarks are displaced from one developmental stage to another will reflect developmental differences.

Our results could probably be improved through the use of more sources of data on the locations of proveins and compartment boundaries early in development. For example, staining of the Wingless expression domain in the larval and pupal wings would allow adding new landmarks by visualization of the hinge/blade boundary in the larval and pupal wings, as well as the anterior and posterior proximal margins (Kolzer et al., 2003). In addition, staining L2 vein domain with antibodies against p-Mad or Srf (Cordero et al., 2007) would also add a new landmark and improve wing shape measurement.

The developmental stage at which differences between the control (*yw*) and the two mutant genotype (*ds* and *shf2*) varied. In the case of *shf2*, the major pattern of variation between the adult *shf2* wings and the *yw* adult wings was evident at the earliest stage studied. Larval, pupal and adult *shf2* wings all had reduced spacing between veins L3-L4 and reduced area compared to *yw*. This suggests that the developmental processes causing this pattern of variation act early in larval development. Previous studies of the *shf2* allele are consistent with our findings (Glise et al. 2005; Gorfinkiel et al. 2005). The *shf* gene codes for a protein involved in the stabilization and diffusion of Hedgehog (Hh) in the larval wing disc. The boundary of Hh signaling in the anterior compartment defines the position of the longitudinal vein L3 along the A-P axis (Blair, 2007). In *shf2*, Shifted fails to properly stabilize Hh, thus shifting posteriorly the Hh signaling boundary and the position of vein L3. In addition, *shf2* wing discs have a reduced expression domain of Dpp, which is a wing growth factor. Thus, the variation in wing shape caused by the *shf2* mutation is due to modification of early larval signaling events.

In contrast, we found that the size and shape difference between *ds* and *yw* have a more complex developmental trajectory. Changes at all the developmental stages we studied contribute to the overall pattern of adult wing shape and size variation between *ds* and *yw*. Some shape differences between *ds* and *yw* appear early during larval development, others during the larval to pupal eversion, and some others during pupal development. For size, the differences appeared during pupal development. As in the case of *shf2*, these findings are consistent with known roles of Dachsous in epithelial morphogenesis, but they also point out to some unknown effects. Dachsous plays an important contribution in orienting cell division during larval development (Baena-López et al., 2005; Mao et al., 2011). In addition, Dachsous mediates cell rearrangements and orientation of cell divisions in response to global tissue stress during pupal development in wing and notum epithelia (Aigouy et al., 2010; Bosveld et al., 2012). Interestingly, our data suggest novel roles of Dachsous in morphogenesis by contributing to tissue shape changes during the larval to pupal transition, as well as to tissue growth during pupal development.

The concordance between the known developmental roles of these well-studied mutations and the differences we observe validates our approach to the quantification of developmental events. It suggests that morphometric studies of shape transformations in genotypes with an unknown developmental basis could provide useful hypotheses about the developmental events involved.

Our work allows us to investigate both the magnitudes of differences in shape and size, and the directions of changes between the developmental stages studied. Consistent with the visually apparent differences in shapes among stages (e.g. Fig. 3), and the relatively dramatic folding and eversion that takes place during pupariation, larval wing shape is more different from pupal wing shape than pupal is from adult wing shape. Differences among individuals with the same genotype at the same developmental stages are noticeably smaller than differences among genotypes. While the differences among stages and genotypes are clear, it is nevertheless apparent that the transformations that each shape undergoes during development is rather similar. This is confirmed by the relatively small angles between developmental trajectories of different genotypes.

## Conclusion

Our approach successfully identified the developmental stage at which variation appears in two cases for which the developmental causes of the variation were known. This suggests that our approach should be useful to study the developmental causes of wing shape variation in cases where we are blind regarding the developmental causes of the variation, as in the case of natural variation.

## Acknowledgments

We thank Jinghua Gui for technical assistance. AMV thanks the Courtier-Orgogozo lab members for stimulating discussions. This research was funded by the Finnish Academy to I. S.-C.

